# Nothing better to do? Environment quality and the evolution of cooperation by partner choice

**DOI:** 10.1101/2020.05.04.076141

**Authors:** Paul Ecoffet, Nicolas Bredeche, Jean-Baptiste André

## Abstract

The effects of partner choice have been documented in a large number of biological systems such as sexual markets, interspecific mutualisms, or human cooperation. There are, however, a number of situations in which one would expect this mechanism to play a role, but where no such effect has ever been demonstrated. This is the case in particular in many intraspecific interactions, such as collective hunts, in non-human animals. Here we use individual-based simulations to solve this apparent paradox. We show that the conditions for partner choice to operate are in fact restrictive. They entail that individuals can compare social opportunities and choose the best. The challenge is that social opportunities are often rare because they necessitate the co-occurrence of (i) at least one available partner, and (ii) a resource to exploit together with this partner. This has three consequences. First, partner choice cannot lead to the evolution of cooperation when resources are scarce, which explains that this mechanism could never be observed in many cases of intraspecific cooperation in animals. Second, partner choice can operate when partners constitute in themselves a resource, which is the case in sexual interactions and interspecific mutualisms. Third, partner choice can lead to the evolution of cooperation when individuals live in a rich environment, and/or when they are highly efficient at extracting resources from their environment.

## 1. Introduction

Among the diversity of mechanisms put forward to explain the evolution of cooperation among non-kin, partner choice has been considered over the last twenty years as having probably played a particularly important role (Eshel and Cavalli-Sforza, 1982; Bull and Rice, 1991; West et al., 2007; Schino and Aureli, 2017; Baumard et al., 2013; Noë and Hammerstein, 1994). When individuals can choose among several partners, which they can compare and compete against each other as in an economic market, this generates a selection pressure to cooperate more, in order to appear as a good partner, and attract others’ cooperation (Noë and Hammerstein, 1994).

The effects of partner choice have been well described in many biological systems (Noë et al., 2001). For example, in the interaction between cleaner fishes and their clients, the law of supply and demand determines how the added value of the interaction is shared, following market principles (Bshary and Grutter, 2006). When cleaners are rare, clients tolerate cheating on their part, while they become pickier when cleaners are numerous. The effects of partner choice have been documented in primate grooming, in meta-analyses showing that females groom preferentially those that groom them most and that a positive relationship exists between grooming and agonistic support (Schino, 2007; Schino and Aureli, 2008). In vervet monkeys, experiments have shown that individuals groom others in exchange for access to food, and do so for longer periods when fewer partners are available (Fruteau et al., 2009). The effects of partner choice have also been documented in humans, where it has been shown that the need to attract social partners is a major driver of cooperation (Barclay and Willer, 2007; Baumard et al., 2013; Barclay and van Vugt, 2015; Debove et al., 2015; Barclay, 2016). Besides, beyond cooperation partner choice also plays a decisive role in mating, leading to the evolution of secondary sexual characteristics, nuptial gifts, and assortative matching (Zahavi, 1975; Andersson and Simmons, 2006; Hammerstein and Noë, 2016).

On the other hand, there are many other biological situations in which one would typically expect partner choice to also play an important role, but where no such effect has ever been demonstrated. These include most intraspecific collective actions in non-human animals. The lack of partner choice is particularly salient in collective hunts such as colobus hunting in chimpanzees, or pack hunting in carnivores. No empirical evidence in these species suggests that individuals cooperate for reasons related to partner choice, either to attract partners or to be accepted by them in their hunts. On the contrary, the majority of available data are consistent with the more parsimonious explanation that individuals are merely doing what is in their immediate best interest at any given time (Packer, 1986; Packer and Ruttan, 1988; Melis et al., 2008, 2011). In particular, if cooperation in collective hunts were driven in part by the need to appear as a good partner, individuals would be expected to willingly share the product of their hunts in a way that depends on everyone’s actual engagement, to encourage participation in other hunts in the future. However, such voluntary and conditional sharing has never been documented in animal collective hunts (Melis et al., 2011). In evolutionary terms, therefore, collective hunting in these species is most likely an instance of *by-product* cooperation, rather than an instance of reciprocal cooperation based on partner choice. This lack of observation is all the more surprising given that, in similar collective actions, human behaviours are demonstrably driven by the need to appear as a good partner (Alvard and Nolin, 2002; Baumard et al., 2013). One may therefore wonder why the same effects did not produce the same consequences in other species.

Such a lack of observation could always be the consequence of the methodological difficulty in empirically proving the existence of partner choice, and more generally of conditional cooperation, outside humans (McElreath et al., 2003; Raihani and Bshary, 2011). However, we would like to suggest an alternative here, namely that there is in fact a strong constraint impeding partner choice in many situations.

Partner choice requires that individuals can compare and choose among several opportunities for cooperation. In some cases, *partners* themselves are opportunities for cooperation and partner choice then only requires that partners are many and accessible. This is the case, for instance, in mating markets, or most instances of interspecific mutualism.

In other cases, however, finding an opportunity for cooperation requires more than just finding a partner. This is what happens when cooperation consists of several individuals working together to exploit environmental resources. In this case, a cooperation opportunity requires both a partner(s) and a resource, which imposes an additional constraint limiting the scope of partner choice. When resources are scarce, there are always few options to compare, and partner choice cannot operate. This could explain the lack of cooperation, beyond by-product cooperation, in many instances of collective actions in the wild despite the availability of potential partners.

To our knowledge, all models published so far on the evolution of cooperation by partner choice focus on situations where finding a partner is sufficient to create an opportunity to cooperate. In this case, they show that partner choice can drive the evolution of cooperation in a relatively wide range of circumstances (Noë and Hammerstein, 1994; Johnstone and Bshary, 2008; Aktipis, 2004; McNamara et al., 2008; Aktipis, 2011; Barclay, 2011; André and Baumard, 2011a,b; Campennì and Schino, 2014; Debove et al., 2015, 2017; Geoffroy et al., 2019). In this paper, we wish to examine what happens on the contrary when re-source availability constitutes a constraint on the operation of partner choice. To do so, we simulate the evolution of agents placed in an environment containing resources that can be exploited collectively.

We show that, in a low-resource environment, and even if there are plenty of partners, partner choice is not able to drive the evolution of cooperation as individuals cannot pit the few cooperation opportunities against each other. What is more, we also show that the number of potential partners harms the evolution of cooperation when patches are scarce. When potential partners are numerous relative to the number of patches available, there are always too many individuals on any given resource as individuals have nothing else to do anyway. Hence, there is no point in trying to attract partners but on the contrary there are benefits in trying to limit their number. Partner choice is thus only effective when the number of available partners lies within a precise range of values, all the narrower as the availability of patches is low.

We believe that this constraint plays a central role in explaining that, in many species, although individuals do participate in collective actions, sometimes finely coordinating their behaviour with that of others, they do not actually seek to cooperate beyond what is in their immediate personal interest.

## 2. Methods

We built an individual-based model with a population of *N* individuals living in a non-spatialized environment consisting of *ω* different patches on which resources are located. All patches are at the same distance from each other. Each patch can host an unlimited number of individuals who gain payoff units by playing a modified version of the prisoner’s dilemma with all the other individuals present on the same patch (see details on the payoff function below). A simulation is composed of *G* generations, and each generation lasts *T* time steps during which individuals gather payoff units. At the end of the *T* time steps, individuals reproduce in proportion to their total payoff and die. During a time step, every individual is considered one by one in random order. When its turn comes, an individual evaluates each of the *ω* patches of the environment, including the patch where it is currently located, assigns each a score (details in section 2.1), and then moves toward the patch with the highest score, or stays on its current patch if that is the one with the highest score. We assume that individuals can freely choose which patch they go to, and that they cannot be forcibly removed from a patch. When an individual moves to a patch different from its current one, it incurs a cost *c_m_*. Once every individual has taken its decision, individuals express their cooperation strategy on their local patch, and they collect a payoff that depends on their cooperation strategy, their partners’ strategies, and the number of individuals present on the patch. Patches can disappear at every time step, with a probability *d*, and are then immediately replaced by an empty patch. Table 1 sums up the parameters of the model.

**Table 1:** Parameters of the simulation.

| Parameter | Description | Value |
| --- | --- | --- |
| **Environment** |  |  |
| $M$ | Demographic population size | 100 |
| $d$ | Probability of the disappearance of patches, per time step | 1/1 000 |
| $T$ | Number of time steps per generation | 1 000 |
| $c_m$ | Cost of moving to another patch | 0 * |
| $N$ | Social population size | variable |
| **Payoff** |  |  |
| $a$ | Immediate personal benefit of cooperation | 5 |
| $b$ | Social benefit of cooperation | 5 |
| $\hat{n}$ | Optimal number of individuals per patch | variable |
| $\sigma$ | Tolerance to variations in the number of individuals per patch | variable |
| **Evolution** |  |  |
| $G$ | Number of generations | 1 500 |
| $\mu$ | Probability of mutation per gene per generation | 0.01 |

\* Simulation results with a non-zero cost are available in the supplementary material

### 2.1. The decision-making mechanisms

The individuals’ strategy in this environment consists of two separate decisions.

On the one hand, the individual must evaluate the different patches available and assign a score to each. This decision is made by an artificial neural network, called the “patch ranking” network. For each patch, this neural network has the following input information: (i) the number of other individuals already present on the patch, (ii) the average level of cooperativeness expressed by these individuals in the last time step, (iii) the level of cooperation that the focal individual would express should it join this patch, and (iv) a boolean that indicates whether the individual would have to move in space to join this patch (i.e. this boolean distinguishes the patch where the individual is currently located from all other patches). This patch ranking network allows the individual to implement partner choice behaviour by choosing where (and thus with whom) it wishes to interact.

On the other hand, the individual must decide on a level of cooperativeness once it is on a patch.

This decision is made by another artificial neural network called the “cooperation” network. As an input, this neural network only has the number of other individuals present on the same patch as the focal. We assume that the agent cannot modulate its cooperation level as a function of others’ cooperation level. This assumption is meant to exclude the possibility that partner control strategies may evolve and allows us to focus only on the effect of partner choice (Noë and Hammerstein, 1994).

The details of the architecture of the neural networks are available in the supplementary materials. The connection weights of both networks constitute the genome of each agent. They evolve by natural selection as exposed in section 2.4.

### 2.2. Phenotypic variability of cooperation

Each individual *i* present on a patch invests a given amount *x_i_* into cooperation — where *x_i_* is decided by the individual’s cooperation network. However, as is now well established in the literature, selective pressures in favour of conditional cooperation stem from the presence of some variability in partners’ cooperative behaviour (see McNamara and Leimar, 2010). In order to capture the effect of variability in the simplest possible way, here we consider the effect of phenotypic variance in the expression of individuals’ genes.

At each generation of our simulations, each individual is subject to the effect of a *phenotypic noise* that modifies its cooperation level. If 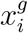 is the cooperation level decided by the cooperation network of individual *i*, then the actual cooperation level played by the individual is 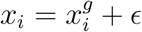, where *ϵ* is drawn randomly as follows. The interval [−1,1] is uniformly split in *N* values, and every individual gets one value of *ϵ* chosen among these *N* values without replacement.

### 2.3. The payoff function

Individuals present on the same patch play a modified version of the n-player prisoner’s dilemma. Consider a focal individual *i* playing *x_i_*, in a patch on which there are *n* – 1 other individuals whose average level of investment is 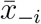. The payoff of individual *i* is given by

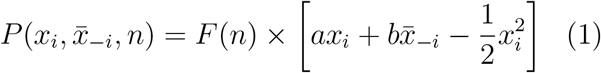

where *a* represents the immediate, self-interested, benefit of the interaction, and *b* represents the social benefit of cooperation from others. When *a* = 0, the game is a prisoner’s dilemma. In our simulations, however, we always choose *a* > 0 which entails that cooperation always has a slight immediate benefit. This assumption is necessary to avoid a bootstrapping problem in the joint evolution of cooperation and partner choice, which is an important issue but not the object of the present paper (see André, 2014 for a similar issue in the case of partner control). Finally, the individual cost of cooperation is assumed to be a quadratic function of the agent’s cooperation level. We make this choice so that cooperation has diminishing net returns and evolution can thus lead to an intermediate level of cooperation in equilibrium.

The function *F*(*n*) is meant to capture the fact that there is an optimal number of individuals exploiting a patch and is given by

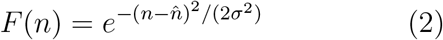

where 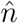 is the optimal number of individuals per patch and *σ* measures the tolerance to variations in the number of individuals per patch (i.e., *σ*^-1^ measures the strength of the penalty that stems from being a suboptimal number of individuals on the same patch). When tolerance *σ* is very low, individuals get a benefit almost only when they are exactly the optimal number 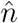 on a patch. On the other hand, when tolerance is very large, there is almost no penalty for being too many, or too few, partners per patch.

With this payoff function, in the absence of partner choice, the evolutionarily stable strategy is to invest *x_ESS_* = *a*, whereas the “socially optimal” cooperativeness level, that is the level that would maximise the average payoff of individuals on the patch, is 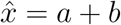.

### 2.4. The evolutionary algorithm

Each individual has a genome composed of the weights of its two neural networks, which makes a total of 84 genes *g* = (*g*_1_,…,*g*_84_) with *g_i_* ∈] – 10,10[. We consider a population of fixed size *M*. The first generation is composed of *M* individuals with random genes for the neural network weights, drawn uniformly in [– 1,1]. We then use a fitness proportionate evolutionary algorithm to simulate evolution. After the *T* time steps of a generation have taken place, individuals all reproduce and die. A new population of *M* individuals is built out of the previous generation by sampling randomly among the *M* parents in proportion to their cumulated payoff, according to a Wright-Fisher process.

A mutation operator is applied to each offspring. Every gene of every offspring has a probability *μ* to mutate and a probability 1 – *μ* to stay unchanged. If a gene *g_i_*, with value *υ_i_*, mutates, it has a probability 0.9 to mutate according to a normal distribution and thus reach a new value sampled in 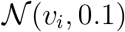 and a probability 0.1 to mutate according to a uniform distribution and thus reach a new value sampled in 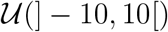.

The evolutionary algorithm is run for *G* generations.

In our analyses, *N* varies, which represents the number of individuals present together in the environment (i.e. the social population size). However, demographic population size *M* must stay constant so as not to alter the relative strength of drift and selection. To do so, we create [*N/M*] parallel environments. The *M* individuals of the demographic population are then randomly assigned so that each environment has exactly *N* individuals. For the last environment to be completed, randomly chosen genetic individuals are duplicated, but their payoff in this environment is then not considered for the calculation of their fitnesses.

## 3. Results

### 3.1. Cooperation cannot evolve when patches are scarce

We simulate the evolution of a population of *N* = 100 individuals for *G* = 1500 generations for different values of the number of resource patches *ω*, but always in a situation where the optimal number of individuals per patch was 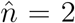. Cooperation only evolved when patches were more abundant than a threshold (Fig. 1a). This can be understood as follows. When resource patches are few, precisely when 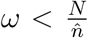, individuals have little cooperation opportunities and there are therefore always more individuals per patch than what would be optimal (in this case, the optimal number of individuals per patch is 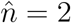). As a result, additional individuals joining a patch are more of a nuisance than a benefit, and there is therefore no benefit in trying to attract partners by appearing cooperative. On the contrary, when the number of available patches is non-limiting, individuals have many social opportunities and it is therefore worth investing in a cooperative behaviour to attract partners. This second situation also captures what happens when individuals themselves constitute a resource.

**Figure 1:**
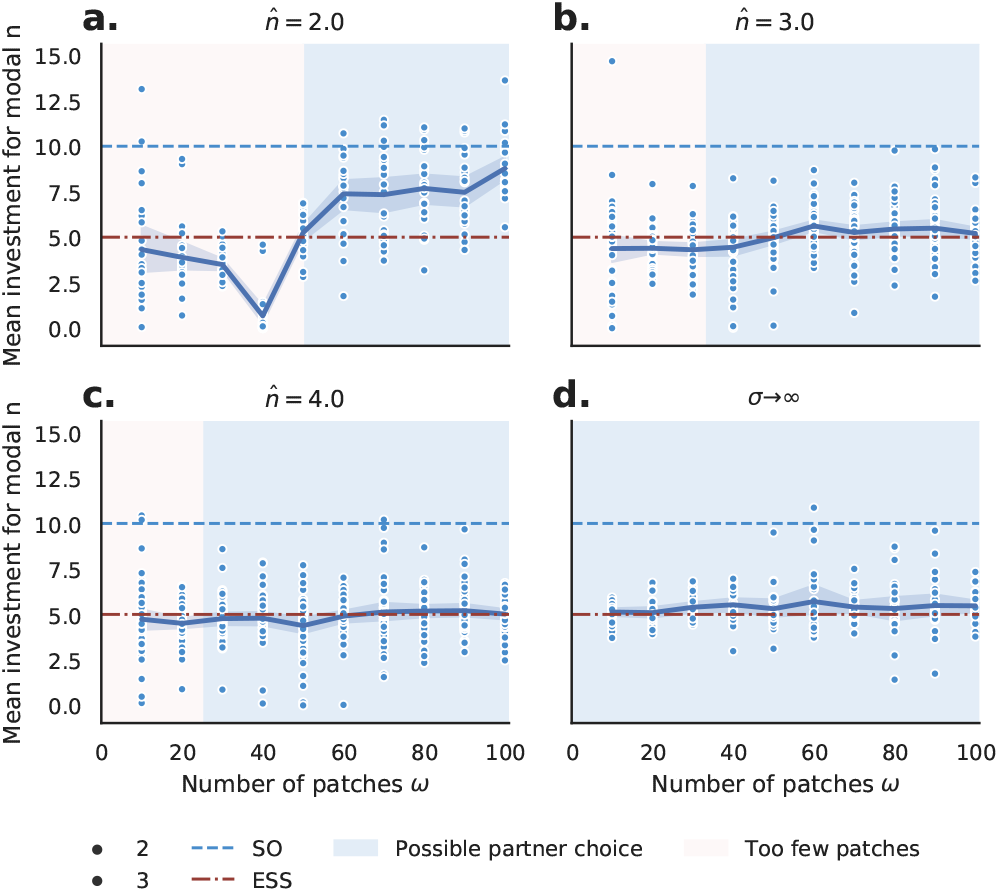
Mean investment in simulation for different numbers of patches *ω* and a fixed population of *N* = 100 individuals. Results after 1500 generations. **a.** When 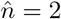, *σ* = 1, individuals invest around or below the optimal selfish investment (ESS) when the number of patches *ω* < 50. They invest above the ESS value and closer to the social optimum value (SO) when *ω* > 50. Cooperative behavior evolves when *ω* ≥ 50. **b-c.** For 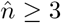, *σ* = 1, individuals invest around the optimal selfish investment value (ESS) regardless of the number of patches *ω*. Cooperative behaviours never evolve. **d.** When *σ* → ∞, there is no constraint on the optimal number of individuals per patch. The individuals’ investment is the ESS value, regardless of the number of patches *ω*. Cooperative behaviours never evolve.

We simulate the evolution of cooperation in situations where the optimal number of individuals per patch, 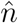, is larger (Figs. 1b, and 1c). Overall, the outcome is even less favourable to cooperation. This can be explained by a “dilution of responsibility” effect. When there are many individuals on a patch, the marginal effect of each individual on overall group performance is very low. Each individual therefore has little effect on the attractiveness of his patch, and this strongly reduces the interest of investing in cooperation to attract partners.

Simulations were run where the tolerance coefficient *σ* varies throughout the conditions. The higher the tolerance on the number of individuals on a patch (i.e. the higher *σ*), the less cooperation is favoured by evolution (results are shown in Fig. 1d in the extreme case where the number of individuals per patch has no impact at all on the payoff, and in supplementary materials in the general case, Fig. A.4). This result can also be understood because there cannot be any benefit in attracting partners when the number of individuals per patch does not matter.

Lastly, we also vary the cost of moving to another patch, *c_m_*. The higher the cost, the less cooperation is favoured, as expected from the literature on partner choice (supplementary materials, Fig. A.5).

Overall, the evolution of cooperation by partner choice can only take place in the restricted conditions where (i) there is an optimal number of individuals per resource patch, (ii) this optimal number is low, and (iii) the number of resource patches in the environment is large.

### 3.2. Cooperation cannot evolve when there are too many partners around

In a second step, we simulate again the evolution of a population of *N* =100 individuals for *G* = 1500 generations in a situation where the optimal number of individuals per patch is 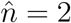, but this time the number of patches is hold constant, *ω* = 20, while varying the actual number of individuals, *N*, present together in the environment (Fig. 2a).

**Figure 2:**
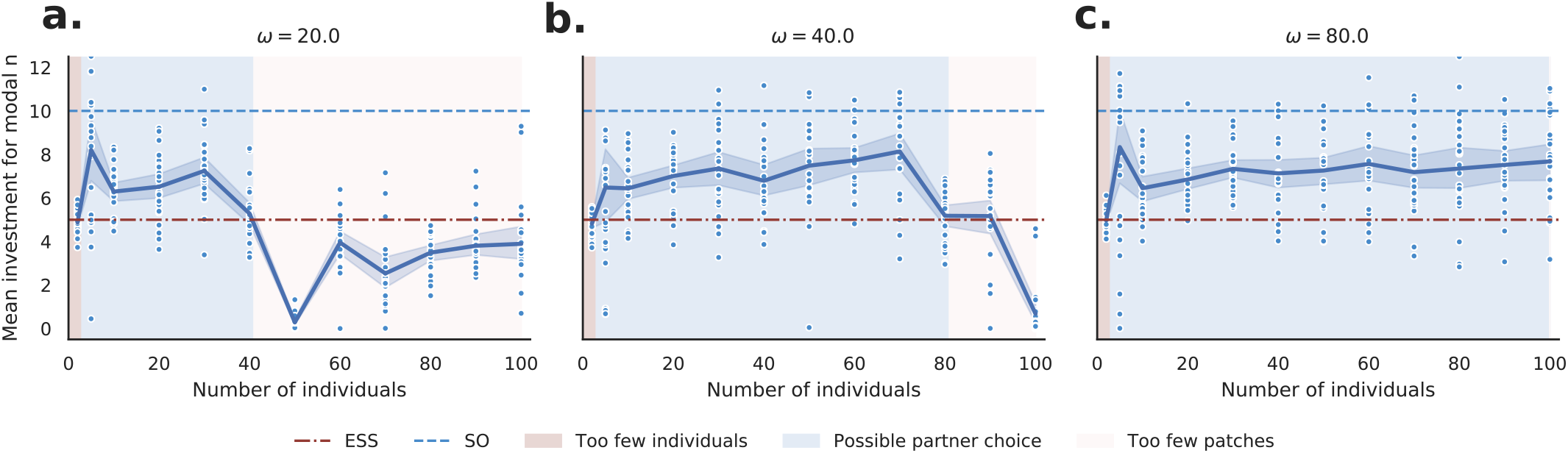
Effect on the population size in the environment with 20, 40 or 80 patches and an optimal number of agents 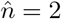 and *σ* = 1. Agents invest above the ESS value and closer to the social optimum (SO) for 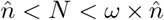. Outside of this range, individuals invest below or about the ESS. Cooperative behaviour is observed only for 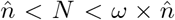. Thee size of this range gets bigger as the number of patches increases.

In this case, cooperation only evolves when the number of individuals in the environment is intermediate. This can be understood as follows. When the number of individuals in the environment, *N*, is too close to the number of individuals 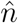 that are needed to exploit at least one patch — or even more so when 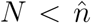, then the number of available partners is limiting. As a result, the actual number of cooperation opportunities from which individuals can choose is very low, partner choice is thus a weak force, and the benefit of investing into cooperation is low. On the other hand, when the number of individuals in the environment *N* is larger than the total number of individuals that can be accommodated on the available patches, that is when 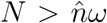, the number of available patches is limiting. The problem is rather that there are always too many individuals on each patch than too few and partner choice is also a weak force. There is, therefore, a range of intermediate population densities, neither too low nor too high, for which cooperation can evolve.

The same simulations are then performed again but with more patches available in the environment (i.e. for larger *ω*, Figs. 2b and 2c). The range of population densities for which cooperation could evolve was then broader. This can again be understood in the above framework. On the one hand, the lower boundary of population density, 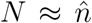, below which the number of individuals is a limiting factor, is unaffected by the number of patches available. On the other hand, the upper boundary of population density, 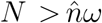, above which the number of patches is a limiting factor, increases with the number of patches, *ω*. As a result, the width of the range of population densities where partner choice is effective increases.

Our model thus highlights two constraints on the evolution of cooperation by partner choice. First, the ratio of the number of individuals per resource patch must be low enough, so that individuals have the opportunity to compare different available patches. Second, the absolute number of individuals in the environment must be high enough, so that individuals have the opportunity to compare different partners.

### 3.3. Analysis of the behaviour of “patch ranking” networks

We analyse the response of the patch ranking networks that were present in our simulations after 1500 generations. To do so, we feed the patch ranking network of each agent with fake patch values (varying the partners’ investment, the focal individual’s investment and the number of individuals present on the patch) and evaluate their response. We adjust the score of each patch to allow for comparisons between individuals and simulations (see details in the supplementary materials).

First, we analyse the networks that evolve when the number of patches in the environment is nonlimiting such that cooperation and partner choice evolve (Figs. 3a and 3b). In this case, we find that the patch ranking networks always prefer to move to patches (i) where there is exactly one partner already present (Fig. 3a) and (ii) where the partner’s level of cooperativeness is the highest (Fig. 3b), which confirms that partner choice, and more generally patch choice, is at work in this condition.

**Figure 3:**
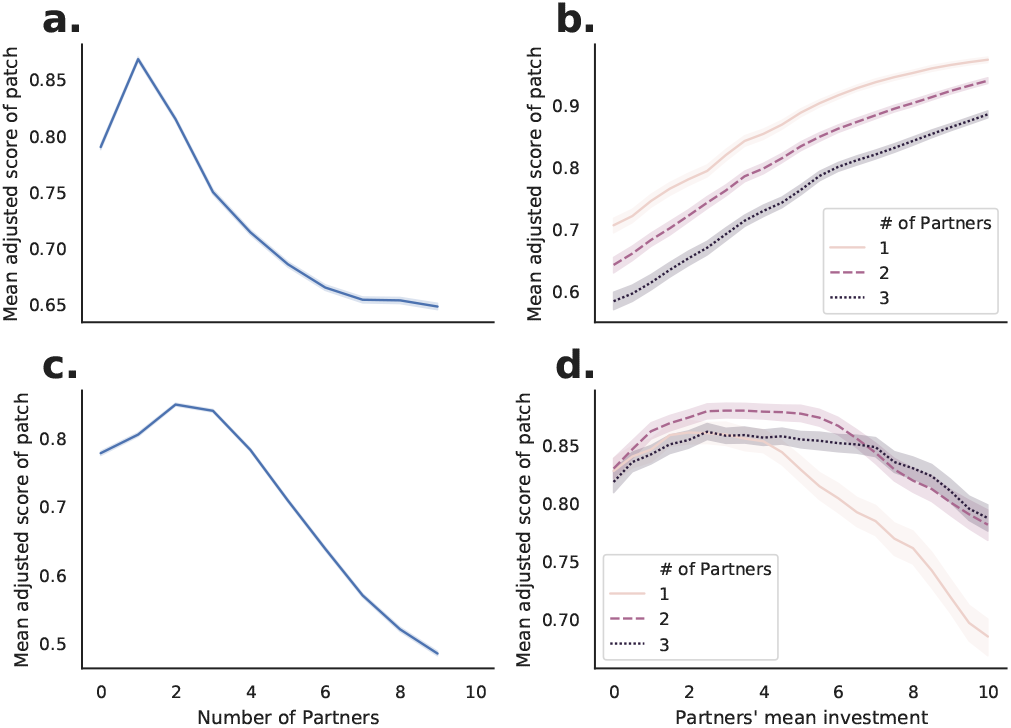
Mean score of patches as evaluated by the patch ranking network of 100 individuals in 24 simulations, with *N* = 100, 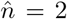, and *σ* = 1. The investment of the focal agent is set to the value it would have invested in the context of the evaluated patch. **a**. Mean patch score as a function of the number of partners already present with *ω* = 80. Mean individuals’ score is the highest for patches with a partner already present (therefore with two individual including themselves). The mean score decreases as the number of partners on the patch increases, or when the individual is alone on the patch, i.e. when the number of partners is zero. **b**. Mean patch score as a function of partners’ mean investment with *ω* = 80. Mean patch score increases as the partners’ mean investment increases. Individuals always prefer the most cooperative partner available. This is characteristic of a partner choice response. **c**. Mean patch score as a function of the number of partners already present with *ω* = 20. The patches scores are the highest when there is already two or three individuals on a patch. Individuals prefer patches where there are already two or three individuals on the patch, even though the optimal number of individuals on a patch is 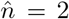. **d**. Mean patch rank as a function of partners’ mean investment with *ω* = 20. Mean patch scores are the highest around the ESS value *x_ESS_* = 5 or the score are mostly flat regardless of the partners’ mean investment. Individuals prefer patches where partners invest around the value *x_ESS_* = 5 or do not have a clear preference about their partners’ investment. Patch choice and therefore partner choice mechanisms have not developed.

Second, we analyse the networks that evolve when the number of patches in the environment is highly limiting such that cooperation and partner choice cannot evolve (Figs. 3c and 3d). In this case, the patch ranking networks always prefer to move to patches (i) where there are already two or three partners present, (Fig. 3c) and (ii) where the partners’ level of cooperation is close to the selfish optimum level of cooperation (Fig. 3d). This confirms that, in this case, partner choice is not a driving force able to lead to the rise of cooperation.

## 4. Discussion

Partner choice can lead to the evolution of cooperation when individuals can compare several opportunities for social interaction and choose the most advantageous ones. In this article, we have shown that the conditions for this to happen are, however, quite restrictive. They entail that individuals truly have access to a range of social opportunities. However, in many cases, social opportunities are rare because they necessitate the co-occurrence of two things at the same time: (i) at least one available partner, and (ii) an exploitable resource or, more generally, “something to do” with that partner. In this article, we have used individual-centred simulations to study the consequences of this constraint on the evolution of cooperation by partner choice. We have obtained the following results.

First, partner choice cannot lead to the evolution of cooperation when resources are scarce, and therefore opportunities for cooperation are rare. This explains why, in many species, social interactions show no evidence of cooperation beyond immediate self-interest (Scheel and Packer, 1991; Bullinger et al., 2011; Melis et al., 2011). Even when individuals engage in collective actions, for example when they hunt collectively, others have so few alternative opportunities anyway that there is no need to seek to draw them into the collective actions. They will come anyway, for want of anything better to do. Even worse than that, as opportunities for cooperation are rare, not only are there always enough partners in each collective action without it being necessary to actively attract them. In fact the opposite is true: There are always *too many* individuals participating in each cooperation endeavour (see Fig. 2). This has been documented for instance in pack hunting in Lions, where Packer showed that lionesses often hunt in groups that are too large compared to what would be optimal (Packer et al., 1990). In such a case, the average gain per individual in a collective action is reduced and not increased by the participation of others, and there is therefore no selection to attract partners but rather a selection to push them away at the time of sharing.

Second, partner choice can lead to the evolution of cooperation when partners constitute in themselves resources. There is, in this case, no further requirement for a social opportunity than the need to find a partner. This occurs, for instance, in sexual markets, or in the many instances of interspecific mutualisms, where the other individual alone constitutes an opportunity to cooperate. It is therefore understandable that partner choice plays a particularly important role in these two types of interactions (Bshary and Grutter, 2002; Andersson and Simmons, 2006; Schino and Aureli, 2008).

Third, partner choice can lead to the evolution of cooperation when the environment is rich or, said differently, when individuals are efficient at finding opportunities for cooperation in their environment. Living in an environment rich in opportunities, and/or having skills that increase the effective number of opportunities one can exploit, brings with it the possibility of *choosing* between different opportunities. This puts greater pressure on individuals, who are then competing to attract partners on their own opportunity, rather than on another, and thus selects for cooperation beyond immediate self-interest.

Note that these results hinge on the assumption that individuals cannot be forcibly removed from a patch. As a result, the only way to choose one’s partners is to walk away, and thus give up a resource patch entirely. If certain physically stronger individuals had the capacity to forcibly displace others, they would be able to choose their partners even when resources are scarce. However, to study the consequences of this possibility would require an explicit consideration of individual variability in physical strength, which we did not consider in this paper.

Finally, our results could suggest a potential relationship between the evolution of cooperation on the one hand, and the evolution of cognitive abilities to more efficiently extract resources from the environment on the other. The possibility that there is an evolutionary relationship between cooperativeness, or more generally sociality, and cognitive capacities has long been discussed in the literature, and several hypotheses have been proposed to explain it (e.g. (dos Santos and West, 2018; Dunbar and Shultz, 2007)). These hypotheses, however, are all about the joint evolution of sociality with cognitive capacities that are *specifically* dedicated to social life itself. The present results suggest that cognitive abilities that have nothing to do with cooperation or sociality per se, namely the sheer ability to extract resources from the environment, could also play a role in the evolution of cooperation. This occurs because enhanced cognitive abilities allow transforming and extracting high-value resources from the environment (Kaplan et al., 2000), thereby creating more opportunities for cooperation. As a result, a given environment contains more opportunities for cooperation for individuals with strong cognitive skills, such as human beings, than for the individuals of other species. This then affects the state of the market for cooperation, increasing the amount of competition between alternative social opportunities, thereby selecting for more investment into cooperation to attract partners.

## Conflict of Interest

### Funding

This work was supported by the Agence Nationale pour la Recherche grant ANR-18-CE33-0006 MSR, and by the EUR FrontCog grant ANR-17-EURE-0017.

### Data availability

All data and source code used for the making of this article is available at https://osf.io/p5whz.

### Author’s contributions

PE wrote the code, ran the simulations, did the data analysis and wrote the manuscript. JBA, NB and PE designed the study. JBA and NB coordinated the study and helped draft the manuscript. All authors gave final approval for publication and agree to be held accountable for the work performed therein.

## Appendix A. Supplementary Material

### Appendix A.1. Architecture of the Artificial Neural Networks

The decision module of each individual is composed of two artificial neural networks: The “patch ranking” network and the “cooperation” network.

The “patch ranking” network is a multilayer perceptron with six inputs and one bias, one hidden layer composed of ten neurons and one bias, and one output. The activation function of the network is tanh. The inputs of the neural network are the (i) the investment value of the individual if it comes to the patch, (ii) the mean investment value of its partners if it comes to the patch, the number of partners on the patch split in (ii) units, (iv) tens and (v) hundreds and (vi) the cost of moving to the patch. All inputs are normalised between zero and one. The number of partners is split into units, tens and hundreds to allow better discrimination of small variations in the number of partners on a patch when there are a too large number of individuals in the environment (*N* is large). In total, the number of weights in this neural network is 73.

The “cooperation” network is a multilayer perceptron with two inputs and one bias, one hidden layer composed of three neurons and one bias, and one output. The activation function of the network is tanh. The inputs of the network are the number of partner on the patch split (i) in units and (ii) tens. In total, the number of weights in this neural network is 13.

For both networks, all weights are initialised in the range [–1, 1] and their boundaries are [–10, 10].

### Appendix A.2. Adjusted score computation

To study the response of the patch ranking network between several individuals and simulations, we cannot use the raw output of the network of each individual. Indeed, what matters for the effectiveness of the network is the *relative score* between patches for *one individual*. Therefore, two individuals can have the same patch preference ranking even if they do not use the same scores for each patch.

To take this issue into account, for each individual, we compute the statistical rank of all the 210 fake patches tested according to the scores of the patch ranking network. That is, the patch with the lowest score get the value 1 and the patch with the highest score get the value 210. The statistical rank is computed using the “max” method. In the case of equality — which often happens as the network saturates in its boundary values, the rank given to all the equal patches is the highest rank.

Then, we normalise the ranks for legibility. We divide the rank of each patch for each individual by the highest rank given (210) to get a score between 0 and 1. Finally, we average the score a patch between all individuals and simulations for a given condition, giving us its average normalised score for this condition.

### Appendix A.3. Variation on the tolerance strength

**Figure A.4:**
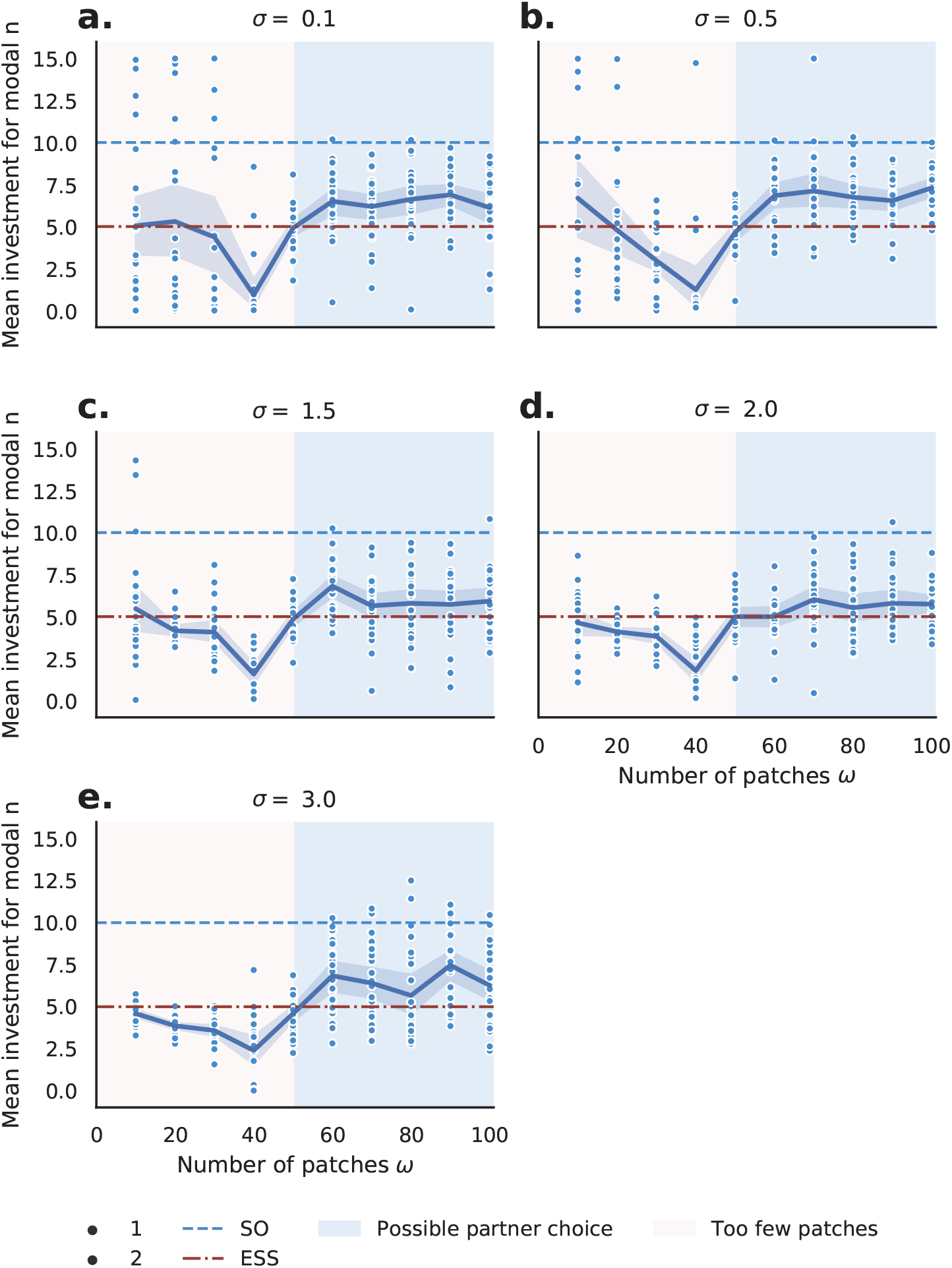
Mean investment in simulations for different numbers of opportunities *ω*, different values of tolerance strengths *σ* and a fixed population of *N* = 100 individuals. Results after 1500 generations. The reference figure when *σ* = 1 is available in Fig. 1a. **a-b.** When the tolerance coefficient is small (i.e. *σ* ≤ 1, see Fig. 1a for *σ* = 1), agents invest above the selfish optimum *x_ESS_* = 5 when *ω* > 50. Agents act cooperatively if the constraint on the environment richness is met. **c-e**. When the tolerance coefficient is too high (i.e. *σ* ≥ 1.5), individuals never invest above the *x_ESS_* value, regardless of the number of patches *ω*. Agents do not cooperate. This is explained by the fact that too many agents (including cheaters) can come on the resource without suffering a cost that has a strong impact on the gains. So there is a dilution effect of responsibility that sets up in the same way as when 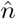 is big.

### Appendix A.4. Effect of the cost of moving

**Figure A.5:**
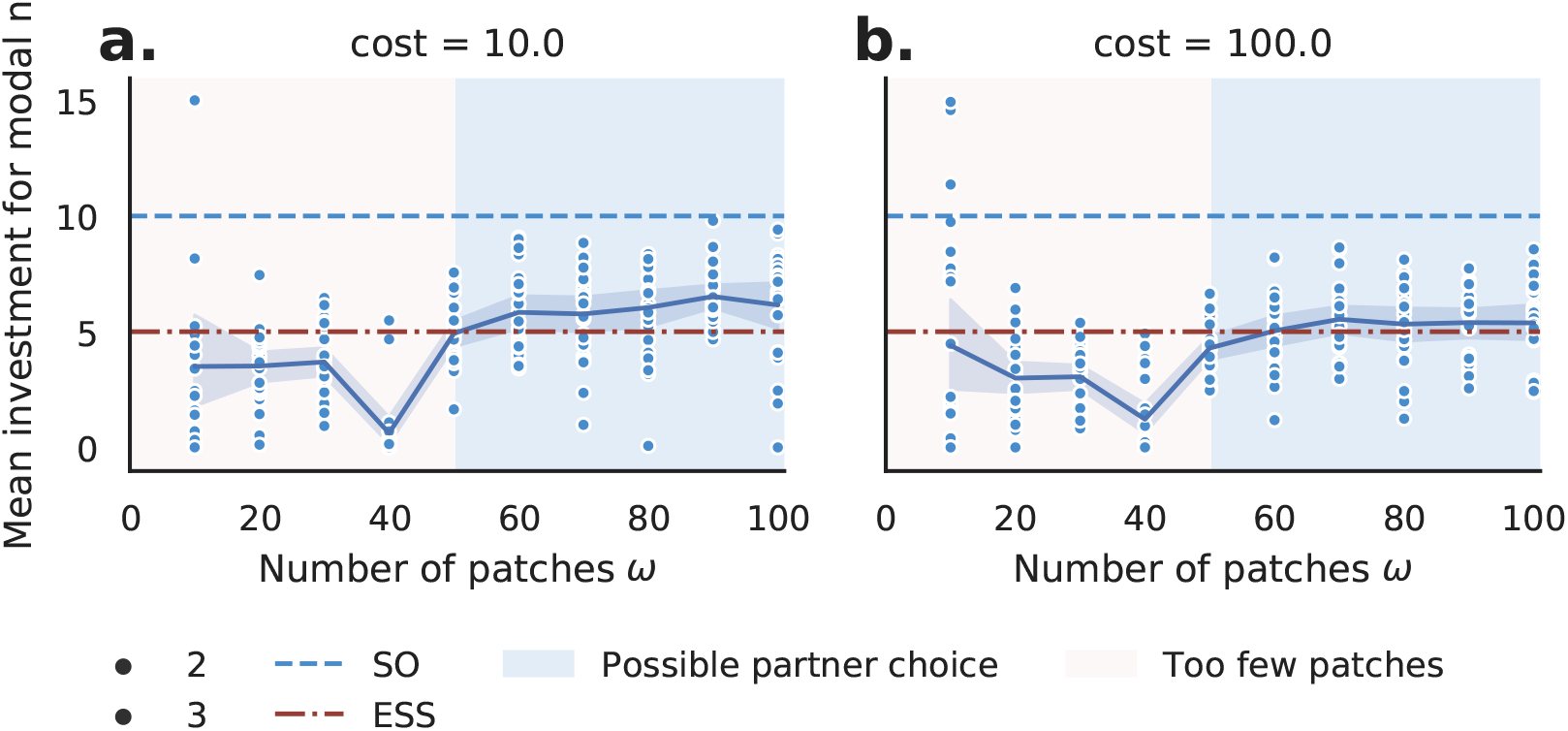
Mean investment in simulations for different numbers of opportunities *ω*, different values of the cost of moving and a fixed population of *N* = 100 individuals. Results after 1500 generations. The reference figure when the cost *c_m_* is 0 is available in Fig. 1a. The greater the cost is, the less cooperative the population is. Increasing the cost of moving increases the cost of partner choice. When the cost is too high, it is of no interest for the agents to cooperate to attract new partners, as if a cheater joins them, it will be too costly for them to leave the opportunity with a defector.

## References

I. Eshel, L. L. Cavalli-Sforza, Assortment of encounters and evolution of cooperativeness, Proceedings of the National Academy of Sciences of the United States of America 79 (1982) 1331–1335. doi:10.1073/pnas.79.4.1331.

J. J. Bull, W. R. Rice, Distinguishing mechanisms for the evolution of co-operation, Journal of Theoretical Biology 149 (1991) 63–74. doi:10.1016/S0022-5193(05)80072-4.

S. A. West, A. S. Griffin, A. Gardner, Evolutionary Explanations for Cooperation, Current Biology 17 (2007) 661–672. doi:10.1016/j.cub.2007.06.004.

G. Schino, F. Aureli, Reciprocity in group-living animals: Partner control versus partner choice, Biological Reviews 92 (2017) 665–672. doi:10.1111/brv.12248.

N. Baumard, J. B. André, D. Sperber, A mutualistic approach to morality: The evolution of fairness by partner choice, Behavioral and Brain Sciences 36 (2013) 59–78. doi:10.1017/S0140525X11002202.

R. Noë, P. Hammerstein, Biological markets: supply and demand determine the effect of partner choice in cooperation, mutualism and mating, Behavioral Ecology and Sociobiology 35 (1994) 1–11. doi:10.1007/BF00167053.

R. Noë, J. A. Van Hooff, P. Hammerstein, Economics in nature: social dilemmas, mate choice and biological markets, Cambridge University Press, 2001.

R. Bshary, A. S. Grutter, Image scoring and cooperation in a cleaner fish mutualism, Nature 441 (2006) 975–978. doi:10.1038/nature04755.

G. Schino, Grooming and agonistic support: a metaanalysis of primate reciprocal altruism, Behavioral Ecology 18 (2007) 115–120. doi:10.1093/beheco/arl045.

G. Schino, F. Aureli, Grooming reciprocation among female primates: a meta-analysis, Biology Letters 4 (2008) 9–11. doi:10.1098/rsbl.2007.0506.

C. Fruteau, B. Voelkl, E. Van Damme, R. Noë, Supply and demand determine the market value of food providers in wild vervet monkeys, Proceedings of the National Academy of Sciences of the United States of America 106 (2009) 12007–12012. doi:10.1073/pnas.0812280106.

P. Barclay, R. Willer, Partner choice creates competitive altruism in humans, Proceedings of the Royal Society B: Biological Sciences 274 (2007) 749–753. doi:10.1098/rspb.2006.0209.

P. Barclay, M. van Vugt, The Evolutionary Psychology of Human Pro-sociality: Adaptations, Byproducts, and Mistakes, Handbook of Prosocial Behavior (2015) 37–60. doi:10.1093/oxfordhb/9780195399813.013.029.

S. Debove, J.-B. Andre, N. Baumard, Partner choice creates fairness in humans, Proceedings of the Royal Society B: Biological Sciences 282 (2015) 20150392. doi:10.1098/rspb.2015.0392.

P. Barclay, Biological markets and the effects of partner choice on cooperation and friendship, Current Opinion in Psychology 7 (2016) 33–38. doi:10.1016/j.copsyc.2015.07.012.

A. Zahavi, Mate selection-A selection for a handicap, Journal of Theoretical Biology 53 (1975) 205–214. doi:10.1016/0022-5193(75)90111-3.

M. Andersson, L. W. Simmons, Sexual selection and mate choice, Trends in Ecology and Evolution 21 (2006) 296–302. doi:10.1016/j.tree.2006.03.015.

P. Hammerstein, R. Noë, Biological trade and markets, Philosophical Transactions of the Royal Society B: Biological Sciences 371 (2016) 20150101. doi:10.1098/rstb.2015.0101.

C. Packer, 19. The Ecology of Sociality in Felids, in: D. I. Rubenstein, R. W. Wrangham (Eds.), Ecological Aspects of Social Evolution, Princeton University Press, Princeton, 1986, pp. 429–451. doi:10.1515/9781400858149.429.

G. Packer, L. Ruttan, The evolution of cooperative hunting, American Naturalist 132 (1988) 159–198. doi:10.1086/284844.

A. P. Melis, B. Hare, M. Tomasello, Do chimpanzees reciprocate received favours?, Animal Behaviour 76 (2008) 951–962. doi:10.1016/j.anbehav.2008.05.014.

A. P. Melis, A. C. Schneider, M. Tomasello, Chimpanzees, Pan troglodytes, share food in the same way after collaborative and individual food acquisition, Animal Behaviour 82 (2011) 485–493. doi:10.1016/j.anbehav.2011.05.024.

M. S. Alvard, D. A. Nolin, Rousseau’s Whale Hunt?, Current Anthropology 43 (2002) 533–559. doi:10.1086/341653.

R. McElreath, T.-H. Clutton-Brock, E. Fehr, D. Fessler, E. Hagen, P. Hammerstein, M. Kosfeld, M. Milinski, J. Silk, J. Tooby, M. Wilson, Group report: The Role of Cognition and Emotion in Cooperation, in: Genetic and Cultural Evolution of Cooperation, 2003. doi:10.4337/9781781006948.00023.

N. J. Raihani, R. Bshary, Resolving the iterated prisoner’s dilemma: theory and reality, Journal of Evolutionary Biology 24 (2011) 1628–39. doi:10.1111/j.1420-9101.2011.02307.x.

R. A. Johnstone, R. Bshary, Mutualism, market effects and partner control, Journal of Evolutionary Biology 21 (2008) 879–888. doi:10.1111/j.1420-9101.2008.01505.x.

C. A. Aktipis, Know when to walk away: Contingent movement and the evolution of cooperation, Journal of Theoretical Biology 231 (2004) 249–260. doi:10.1016/j.jtbi.2004.06.020.

J. M. McNamara, Z. Barta, L. Fromhage, A. I. Houston, The coevolution of choosiness and cooperation, Nature 451 (2008) 189–192. doi:10.1038/nature06455.

C. A. Aktipis, Is cooperation viable in mobile organisms? Simple Walk Away rule favors the evolution of cooperation in groups, Evolution and Human Behav-ior 32 (2011) 263–276. doi:10.1016/j.evolhumbehav.2011.01.002.

P. Barclay, Competitive helping increases with the size of biological markets and invades defection, Journal of Theoretical Biology 281 (2011) 47–55. doi:10.1016/j.jtbi.2011.04.023.

J.-B. André, N. Baumard, The evolution of fairness in a biological market, Evolution 65 (2011a) 1447–1456. doi:10.1111/j.1558-5646.2011.01232.x.

J. B. André, N. Baumard, The evolution of fairness in a biological market, Evolution 65 (2011b) 1447–1456. doi:10.1111/j.1558-5646.2011.01232.x.

M. Campennì, G. Schino, Partner choice promotes cooperation: The two faces of testing with agent-based models, Journal of Theoretical Biology 344 (2014) 49–55. doi:10.1016/j.jtbi.2013.11.019.

S. Debove, N. Baumard, J.-B. André, Evolution of equal division among unequal partners., Evolution 69 (2015) 1–9. doi:10.1111/evo.12583.

S. Debove, N. Baumard, J. B. André, On the evolutionary origins of equity, PLoS ONE 12 (2017) 5–7. doi:10.1371/journal.pone.0173636.

F. Geoffroy, N. Baumard, J.-B. Andre, Why cooperation is not running away, bioRxiv (2019) 316117. doi:10.1101/316117.

J. M. McNamara, O. Leimar, Variation and the response to variation as a basis for successful cooperation, Philosophical Transactions of the Royal Society B: Biological Sciences 365 (2010) 2627–2633. doi:10.1098/rstb.2010.0159.

J. B. André, Mechanistic constraints and the unlikely evolution of reciprocal cooperation, Journal of Evolutionary Biology 27 (2014) 784–795. doi:10.1111/jeb.12351.

D. Scheel, C. Packer, Group hunting behaviour of lions: a search for cooperation, Animal Behaviour 41 (1991) 697–709. doi:10.1016/S0003-3472(05)80907-8.

A. F. Bullinger, A. P. Melis, M. Tomasello, Chimpanzees, Pan troglodytes, prefer individual over collaborative strategies towards goals, Animal Behaviour 82 (2011) 1135–1141. doi:10.1016/j.anbehav.2011.08.008.

C. Packer, D. Scheel, A. E. Pusey, Why lions form groups: food is not enough, American Naturalist 136 (1990) 1–19. doi:10.1086/285079.

R. Bshary, A. S. Grutter, Experimental evidence that partner choice is a driving force in the payoff distribution among cooperators or mutualists: The cleaner fish case, Ecology Letters 5 (2002) 130–136. doi:10.1046/j.1461-0248.2002.00295.x.

M. dos Santos, S. A. West, The coevolution of cooperation and cognition in humans, Proceedings of the Royal Society B: Biological Sciences 285 (2018). doi:10.1098/rspb.2018.0723.

R. I. Dunbar, S. Shultz, Evolution in the social brain, Science 317 (2007) 1344–1347. doi:10.1126/science.1145463.

H. Kaplan, K. Hill, J. Lancaster, A. M. Hurtado, A theory of human life history evolution: diet, intelligence, and longevity, Evolutionary Anthropology 9 (2000) 156–185. doi:10.1002/1520-6505(2000)9:4<156::AID-EVAN5>3.3.CO;2-Z.

